# Prognostic Roles of LncRNA XIST and Its Potential Mechanisms in Human Cancers: A Pan-Cancer Analysis

**DOI:** 10.1101/2021.06.16.448675

**Authors:** Wei Han, Chun-tao Shi, Jun Ma, Qi-xiang Shao, Xiao-jiao Gao, Hao-nan Wang

## Abstract

**Background:** X-inactive specific transcript (XIST), it has been found, is abnormal expression in various neoplasms. This work aims to explore its potential molecular mechanisms and prognostic roles in types of malignancies.

**Methods:** This research comprehensively investigated XIST transcription across cancers from Oncomine, TIMER 2.0 and GEPIA2. Correlations of XIST expression with prognosis, miRNAs, interacting protens, immune infiltrates, checkpoint markers and mutations of tumor-associated genes were also analyzed by public databases.

**Results:** Compared to normal tissues, XIST was lower in BRCA, COAD, LUAD, lymphoma and OV in Oncomine; In TIMER 2.0, XIST was decreased in BRCA, KICH, THCA and UCEC, but increased in KIRC and PRAD; In GEPIA2, XIST was down-regulated in CESC, COAD, OV, READ, STAD, UCEC and UCS. Public databases also showed that XIST was a good indicator of prognosis in BRCA, CESC, COAD, STAD, OV and so on, but a bad one in KIRC, KIRP and so on. From starBase, we found 29 proteins interacting with XIST, and identified 4 miRNAs, including miR-103a-3p, miR-107, miR-130b-3p and miR-96-5p, which might be sponged by XIST in cancers. Furthermore, XIST was linked with immune infiltration, especially T cell CD4+, and was related to over 20 immune checkpoint markers. In addition, XIST was associated with several tumor-associated gene mutations in some cancers.

**Conclusion:** In summary, abnormal expression of XIST influenced prognosis, miRNAs, immune cell infiltration and mutations of tumor-associated genes across cancers, especially BRCA and colorectal cancer. More efforts should be made to detect potential molecular mechanisms of XIST in the carcinogenesis.

## Introduction

Neoplasm has become a global health problem and a leading cause of death worldwide, with about 20 millions new cases and 10 million deaths in 2020 [1]. Although operation, radiotherapy, chemotherapy, targeted therapy and immunotherapy are the main treatment options for carcinomas, patients at the advanced stage still have poor prognosis[2]. Therefore, exploring novel biomarkers is essential for preventing chemoresistance and improving survival rates of cancer patients.

With a length of more than 200 nucleotides, long non-coding ribonucleic acids (LncRNAs) are a highly heterogeneous group of transcripts, and are involved in various biological functions through the epigenetic regulation of genes and the interaction with proteins and RNAs[3,4]. Due to its regulation, abnormal expression of LncRNAs leads to the development and progression of many diseases, especially malignant tumors[5,6]. Therefore, some LncRNAs are considered as novel biomarkers to predict chemoresistance and evaluate prognosis of cancer patients[7,8].

LncRNA X-inactive specific transcript (XIST) locates at Xq13.2 and coats the X chromosome in cis during X chromosome inactivation (XCI)[9]. Emerging investigations report that abnormal expression of XIST takes part in the regulation of various diseases, including solid tumors[10–12]. Previous researches reported that XIST could induce biological behavior and pathological appearance by interacting with several proteins and micro ribonucleic acids (miRNAs), and play key roles in the generation, progression and prognosis of tumors[13,14]. Recent studies have demonstrated that XIST is down-regulated in several cancers and suppresses the progression of tumors, especially breast cancer[15,16]. However, some studies showed that XIST could promote growth and invasion of colorectal cancer cells, and silencing XIST could repress chemoresistance of acute myeloid leukemia[17,18]. These entirely different roles of XIST in carcinomas resulted in discrepancies of its prognostic value among previous researches[19,20].

The tumor microenvironment (TME) is a complex milieu in which immune infiltration can activate or restrain tumor progression and metastasis, which forms the tumor immune microenvironment (TIME)[21,22]. Previous studies discovered that XIST and its downstream regulators were correlated with TIME and PD-L1 expression, and played a critical role in invasion and metastasis of cancers[16,23–25].

Gene mutation is regarded as a crucial factor for malignant transformation and tumor progression[26]. In addition, Gene mutation, such as BRCA1 and BRCA2, affected immune cell infiltration and response to immunotherapy[26]. Abnormal expression of XIST was also related to the mutation of several tumor-associated genes[27,28]. For example, dysregulation of XIST and 53BP1 affected the survival of breast carcinoma patients with BRCA1 mutation[27]. However, whether the mutation of BRCA1 or other genes leads to abnormal expression of XIST is still obscure.

In view of the outstanding contradictions and vagueness above, we conducts a pan-cancer analysis of XIST to elucidate its potential molecular mechanisms and prognostic roles in multiple cancers.

## MATERIALS AND METHODS

### Pan-Cancer analysis of XIST transcription

Oncomine (https://www.oncomine.org/), Tumor Immune Estimation Resource 2.0 (TIMER 2.0, https://timer.cistrome.org/), Gene Expression Profiling Interactive Analysis 2 (GEPIA2, https://gepia2.cancer-pku.cn/) were performed to analyze XIST transcription in multiple cancers. The first database was Oncomine which consisted of more than 80,000 samples of over 20 types of cancers. The threshold in this research was set as the following criteria: gene rank: Top 10%, fold change: 2, and *P* value: 1E-4. The second one was TIMER 2.0 containing more than 10 thousand samples of over 30 cancer types from The Cancer Genome Atlas (TCGA). The next one was GEPIA2 which was a newly developed database for analyzing the RNA sequencing expression data of more than 9,000 tumors and 8,000 normal tissues from the TCGA and the Genotype-Tissue Expression (GTEx) projects.

Names and abbreviations of types of cancers were all listed as follow: ACC: adrenocortical carcinoma; BLCA: bladder urothelial carcinoma; BRCA: breast invasive carcinoma; CESC: cervical squamous cell carcinoma and endocervical adenocarcinoma; CHOL: cholangio carcinoma; COAD: colon adenocarcinoma; DLBC: lymphoid neoplasm diffuse large B-cell lymphoma; ESCA: esophageal carcinoma; GBM: glioblastoma multiforme; HNSC: head and neck squamous cell carcinoma; KICH: kidney chromophobe; KIRC: kidney renal clear cell carcinoma; KIRP: kidney renal papillary cell carcinoma; LAML: acute myeloid leukemia; LGG: brain lower grade glioma; LIHC: liver hepatocellular carcinoma; LUAD: lung adenocarcinoma; LUSC: lung squamous cell carcinoma; MESO: mesothelioma; OV: ovarian serous cystadenocarcinoma; PAAD: pancreatic adenocarcinoma; PCPG: pheochromocytoma and paraganglioma; PRAD: prostate adenocarcinoma; READ: rectum adenocarcinoma; SARC: sarcoma; SKCM: skin cutaneous melanoma; STAD: stomach adenocarcinoma; TGCT: testicular germ cell tumors; THCA: thyroid carcinoma; THYM: thymoma; UCEC: uterine corpus endometrial carcinoma; UCS: uterine carcinosarcoma; and UVM: uveal melanoma.

### Prognosis Analysis

The correlation of XIST expression with prognosis of patients in multiple cancers was analyzed through three public databases, GEPIA2, PrognoScan (http://dna00.bio.kyutech.ac.jp/PrognoScan/index.html) and Kaplan-Meier Plotter (https://kmplot.com/analysis/). Data of PrognoScan mostly come from the Gene Expression Omnibus (GEO) database, and Kaplan-Meier Plotter utilized Affymetrix microarray data from TCGA. We used heat maps, forest plots and KaplanMeier curves to visualize the survival data of cancer patients. Overall survival (OS), disease free survival (DFS), relapse free survival (RFS), disease specific survival (DSS), distant metastasis free survival (DMFS), progression free survival and distant recurrence free survival were main outcomes. The hazard ratio (HR) and 95% confidence interval (CI) were calculated through univariate analysis.

### Exploration of proteins and miRNAs interacting with XIST

We performed starBase (http://starbase.sysu.edu.cn/starbase2/index.php) to identify candidate proteins and miRNAs potentially interacting with XIST by Pearson correlation analyses. In addition, ceRNA network of starBase was conducted to search for possible ceRNAs that could compete with XIST for miRNAs binding. Next, a network of LncRNA - miRNAs - mRNAs was scheduled by the software named GEPHI.

### Tumor Immune Microenvironment Analysis

TIMER 2.0 was utilized to analyzed the relationship of XIST expression with the abundance of infiltrating immune cell types in multiple cancers by Spearman correlation analyses. In addition, we evaluated the correlation of XIST with immune checkpoint markers and immune cell types in each type of cancers by Spearman correlation analysis.

### Pan-Cancer Analysis of representative gene mutation

We performed TIMER 2.0 to analyze the correlation of XIST expression with representative gene mutation by Spearman correlation analyses. 47 representative genes were listed as follow: AKT1, ALK, APC, AR, ARID1A, ASXL1, ATM, BAP1, BARD1, BRAF, BRCA1, BRCA2, BRIP1, CCND1, CDK4, CDKN2A, CHEK2, EGFR, EPCAM, ERBB2, ERBB3, FANCA, FAT1, FBXW7, FGFR1, KDR, KIT, KRAS, MET, MTOR, NBN, NTRK1, NTRK2, NTRK3, PALB2, PIK3CA, PTEN, RAD51C, RAD51D, RB1, RET, ROS1, SMO, STK11, TP53, TP53BP1 and TSC1.

### Statistical Analysis

Differences of levels of XIST transcription between tumors and normal samples were analyzed by *t*-tests. HRs and *P* value were calculated by univariate Cox regression model in PrognoScan, and by log rank test in GEPIA2 and Kaplan-Meier Plotter. Pearson or Spearman correlation analyses were utilized as above. Above all, *P* <0.05 was set as a statistically significant threshold.

## Results

### The mRNA expression level of XIST in multiple cancers

Three databases, Oncomine, TIMER 2.0 and GEPIA2 displayed the transcription levels of XIST in various types of cancers. In Oncomine, compared to normal tissues, XIST expression was obviously lower in several cancers, such as BRCA, COAD, LUAD, lymphoma and OV (Figure 1A). The details of XIST expression in cancers were shown in Table 1. In TIMER 2.0, XIST expression was also decreased in BRCA in comparison of normal tissues (Figure 1B). In addition, XIST was down-regulated in KICH, THCA and UCEC (Figure 1B). However, it seemed that XIST expression was higher in KIRC and PRAD (Figure 1B). As shown in Figure 1C, the expression levels of XIST seemed to be skimble-scamble in different types of human cancers in GEPIA2. Concretely, XIST expression was decreased significantly in CESC, COAD, OV, READ, STAD, UCEC and UCS; but it was up-regulated in five cancers, including ACC, DLBC, LUAD, TGCT and THCA. However, there were no significant differences between other tumors and normal tissues in GEPIA2, including BRCA. On the whole, XIST was abnormally expressed in different cancers, especially in BRCA, OV, COAD and READ.

**Table 1.**
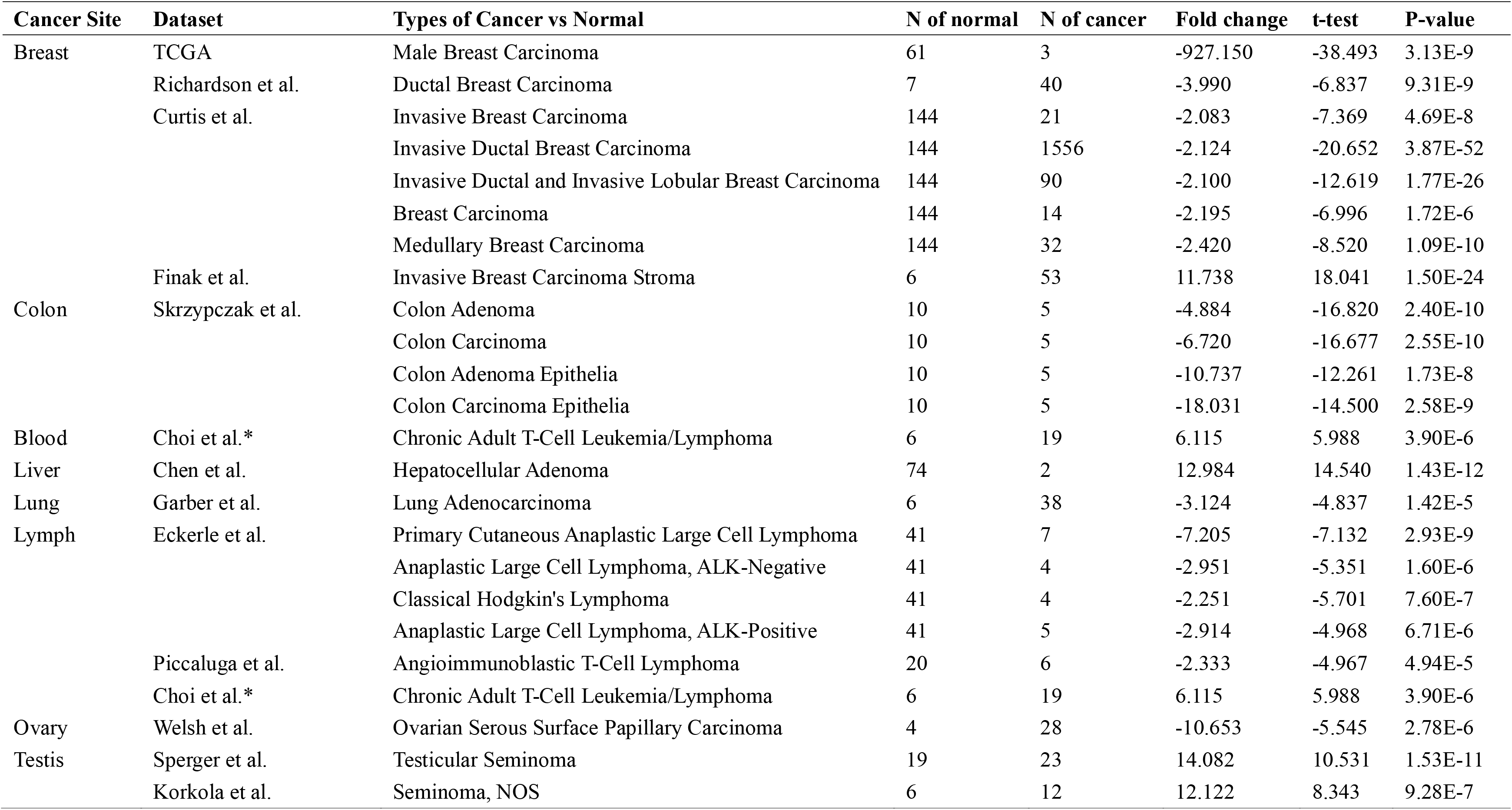
Datasets of XIST transcript across cancers in Oncomine. * Choi et al. analyzed Chronic Adult T-Cell Leukemia/Lymphoma and Chronic Adult T-Cell Leukemia/Lymphoma at the same time.

**Figure 1.**
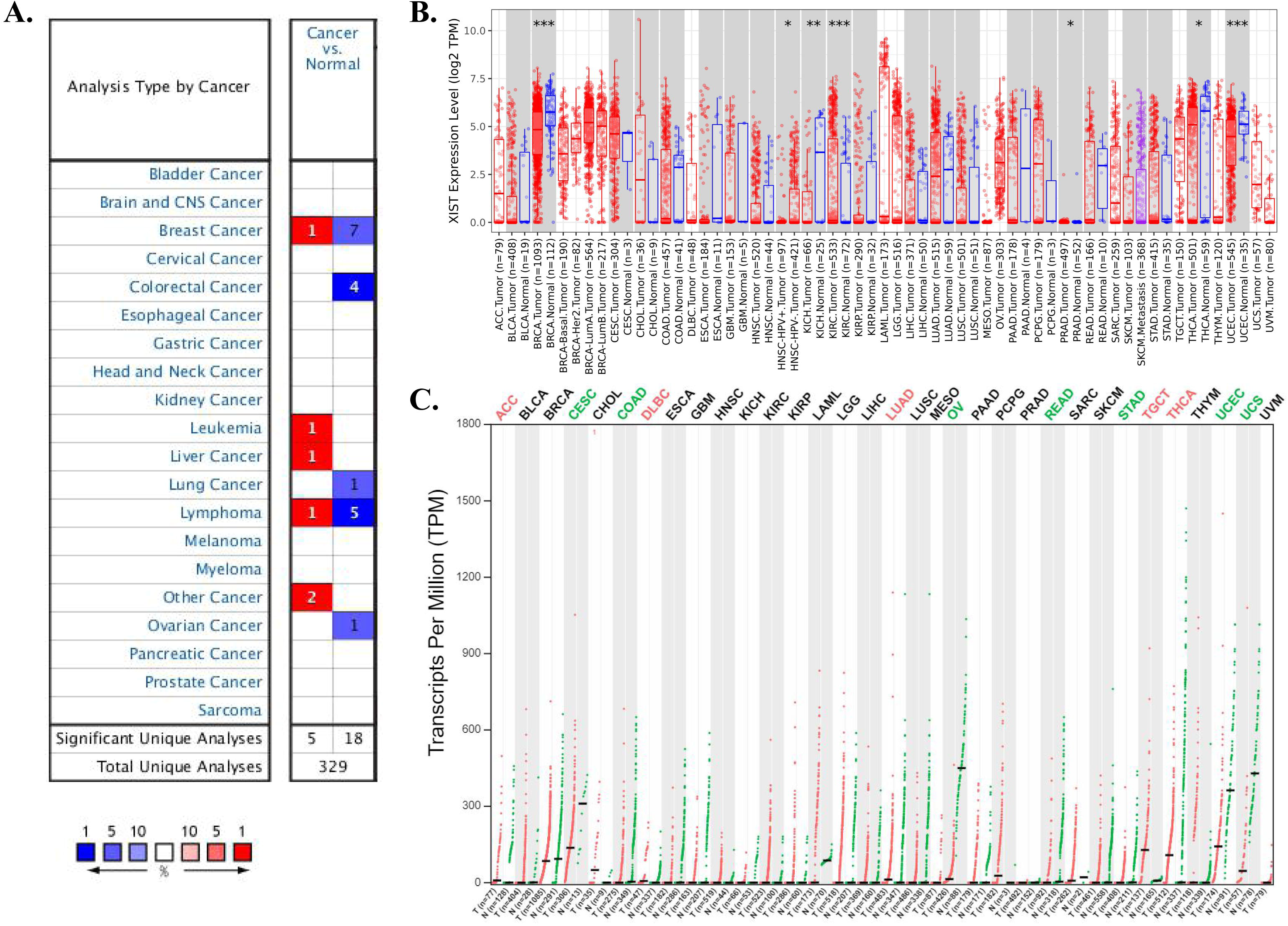
XIST transcript in pan-cancer. **(A)** The transcription levels of XIST in different cancers in Oncomine with exact thresholds (gene rank = 10%, fold change = 2, and *P* < 0.0001). The cell number represented the dataset number with blue for low expression and red for high expression. **(B)** Differential XIST expression between tumors and normal tissues in TIMER 2.0. Red represented tumors and blue represented normal tissues. * *P* < 0.05, ** *P* < 0.01, *** *P* < 0.001. **(C)** Comparison of XIST expression between tumors and normal tissues in GEPIA2. Compared with normal tissues, red represented cancer samples and significantly higher expression in tumors, green represented normal samples and significantly lower expression in tumors, and black represented no significance.

### Prognostic value of XIST in human pan-cancer

Then, we analyzed prognostic role of XIST across cancers in GEPIA2, Kaplan-Meier Plotter and PrognoScan. In GEPIA2, higher expression of XIST indicated longer overall survival rates of patients with CESC and SKCM, but predicted worse prognosis of KIRP (Figure 2A). In addition, XIST expression was positively related to DFS in COAD (Figure 2B). In Kaplan-Meier Plotter, XIST indicated good prognosis of OS in ESCC (*P* = 0.008), PCPG (*P* = 0.0034) and STAD (*P* = 0.047, Figure 3A), and was a favourable factor for RFS in OV (*P* = 0.0059), STAD (*P* = 0.005) and UCEC (*P* = 0.048, Figure 3B). However, XIST was linked to poor outcomes for cancers of HNSC, KIRC, KIRP, LIHC and PAAD. PrognoScan also showed that XIST played an unfavourable prognostic role in several cancers, including BLCA (Figure 4A, OS: Cox *P* = 0.0192), non-small-cell lung cancer (NSCLC, Figure 4L, RFS: Cox *P* = 0.0084) and renal cell carcinoma (RCC, Figure 4N, OS: Cox *P* = 0.0327). But XIST played a protective role in other 6 cancer types, including LAML (Figure 3B, OS: Cox *P* = 0.0324), GBM (Figure 3C, OS: Cox *P* = 0.0285), BRCA (Figure 3D, RFS: Cox *P* = 0.0180; Figure 3E, DSS: Cox *P* = 0.0001; Figure 3F, DFS: Cox *P* = 0.0056), COAD (Figure 3H, OS: Cox *P* = 0.0229; Figure 3I, DSS: Cox *P* = 0.0228; Figure 3J, DFS: Cox *P* = 0.0343), LUAD (Figure 3K, OS: Cox *P* = 0.0004) and OV (Figure 3M, OS: Cox *P* = 0.0074). However, it seemed that XIST was not a favourable factor for DMFS in BRCA (Figure 3G, Cox *P* = 0.0231). The details of survival analyses in PrognoScan were listed in Table 2.

**Table 2.**
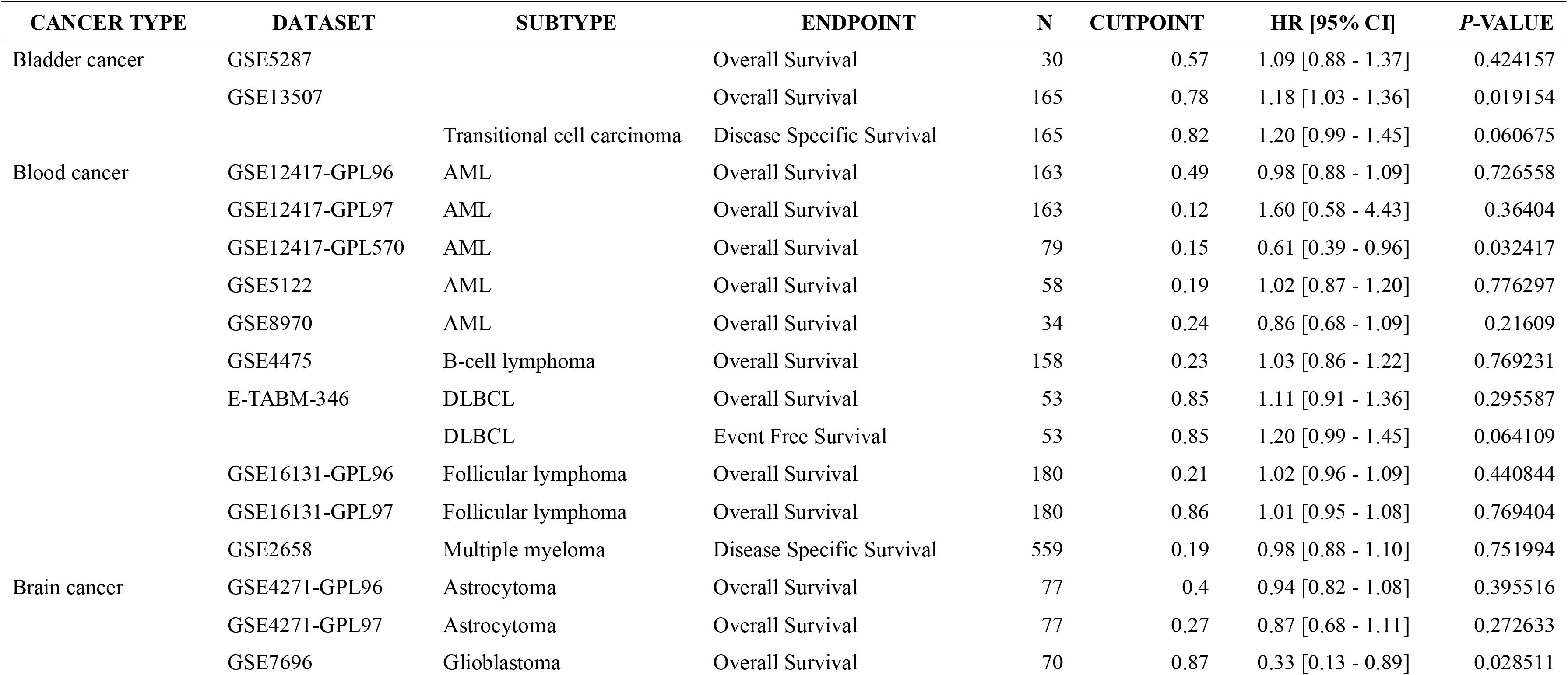
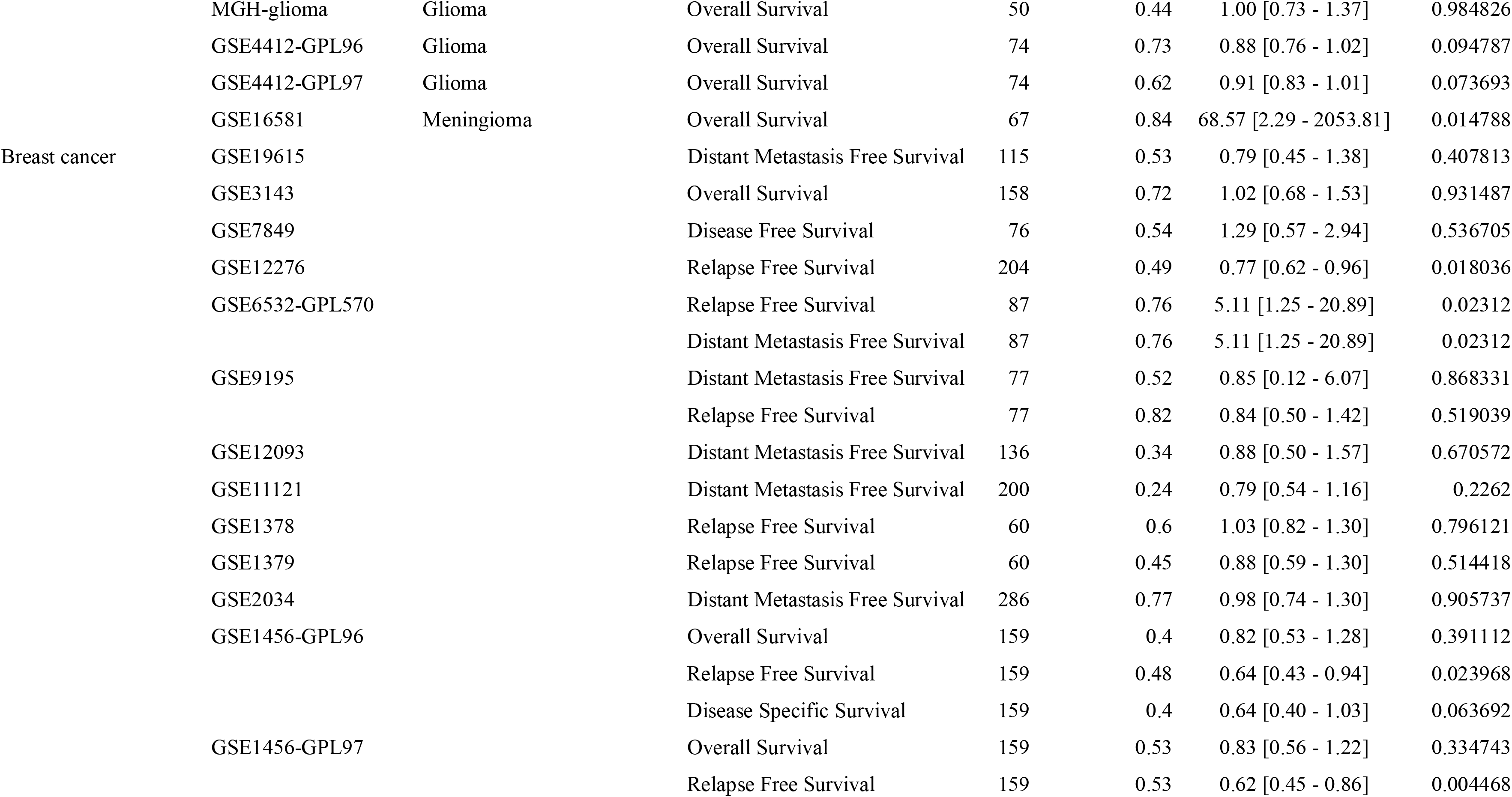
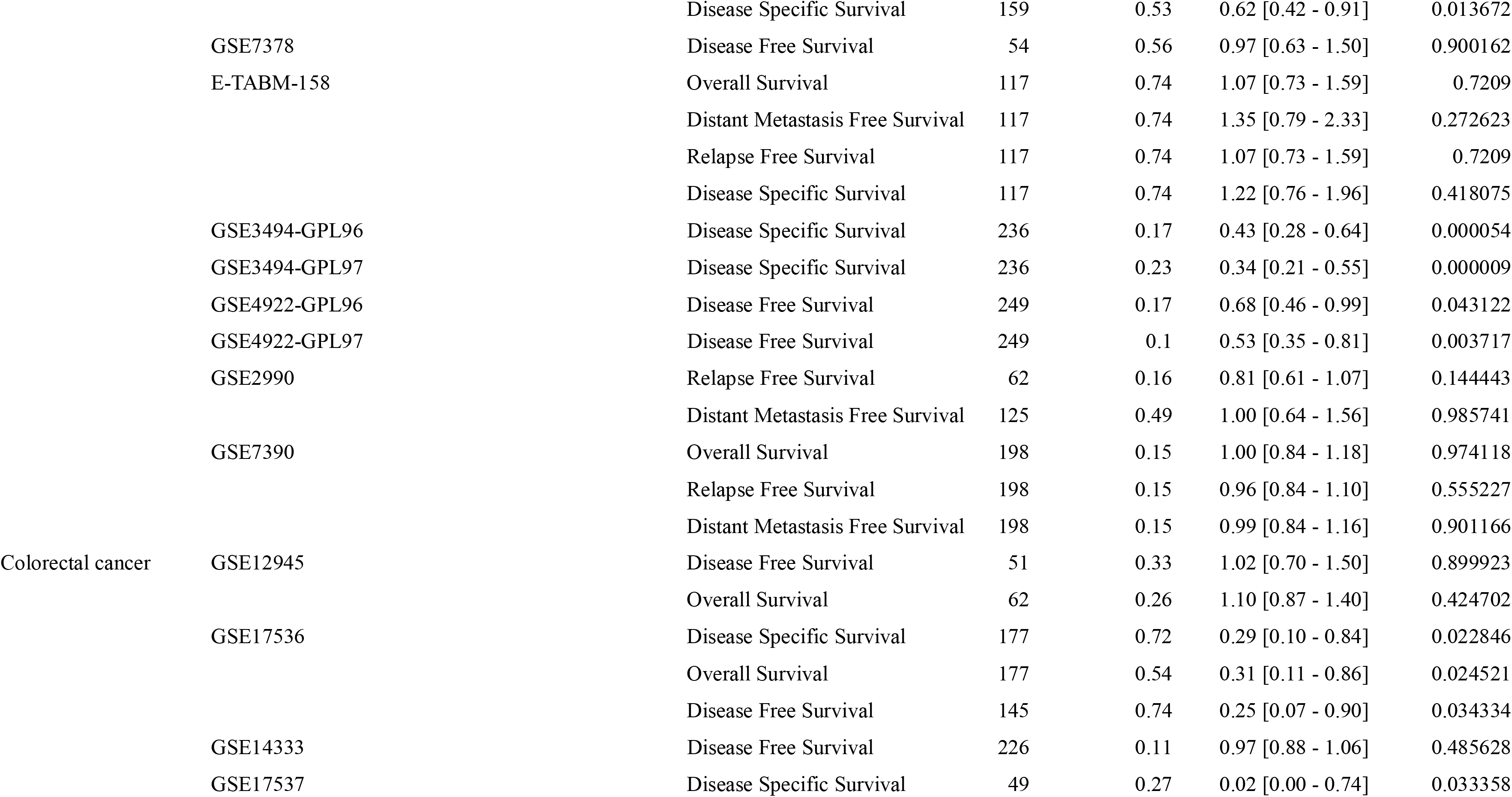
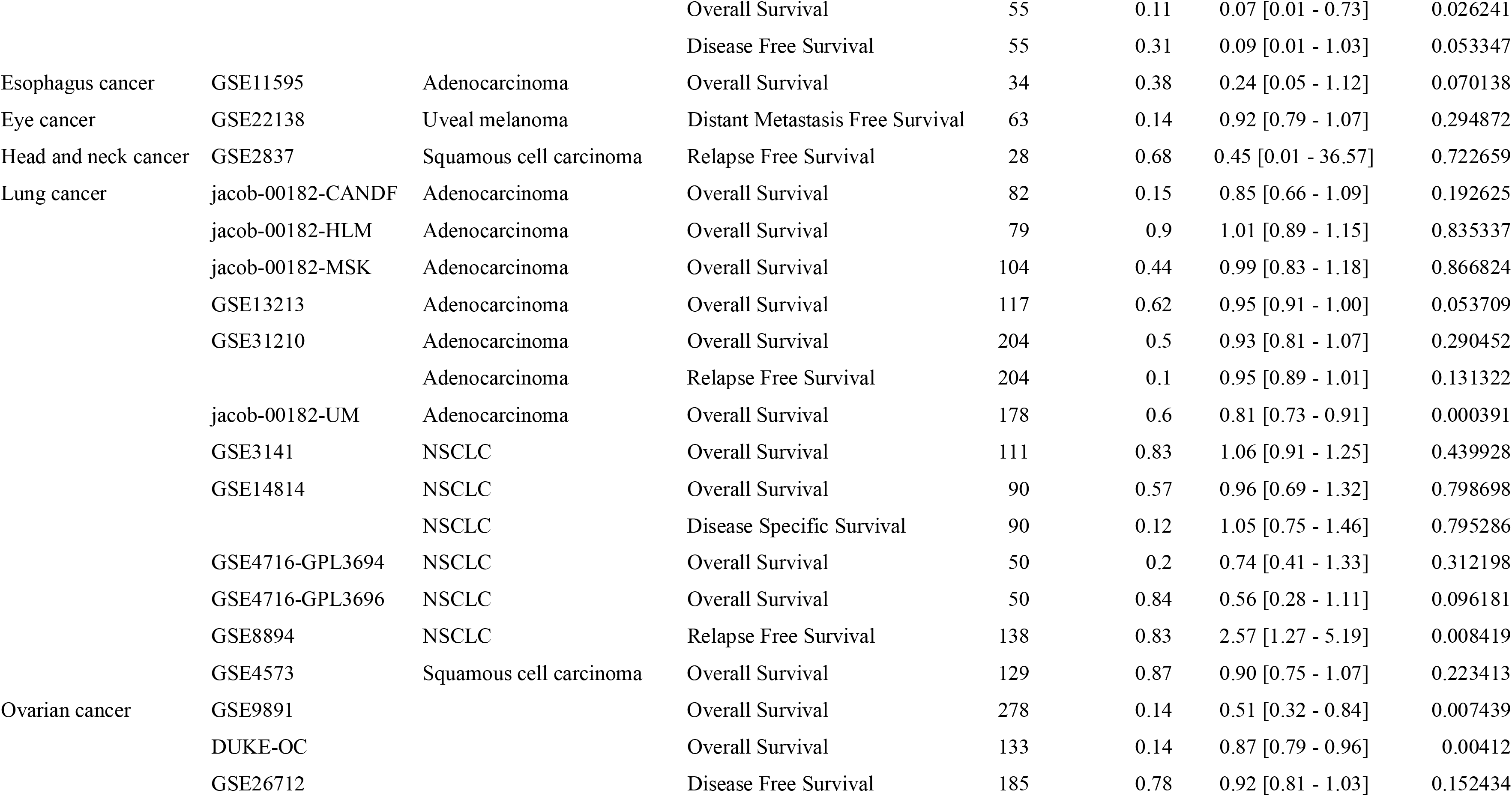
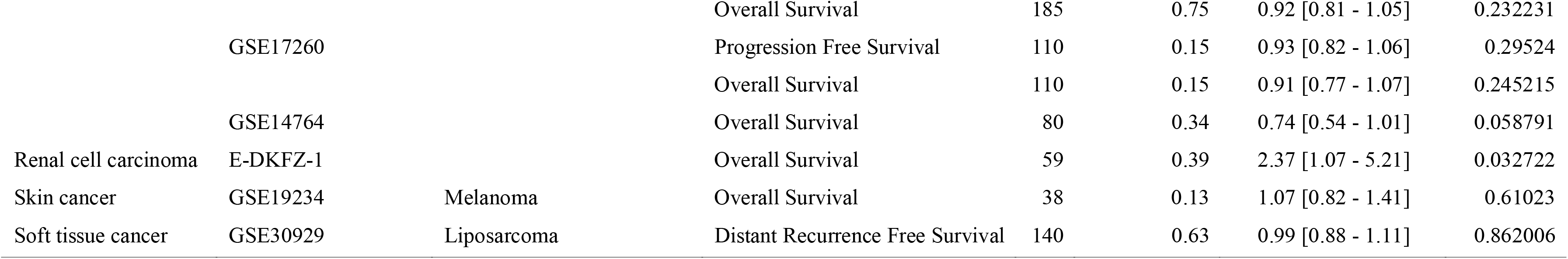
Prognosis analyses of XIST expression across cancers by univariate Cox regression model in PrognoScan.

**Figure 2.**
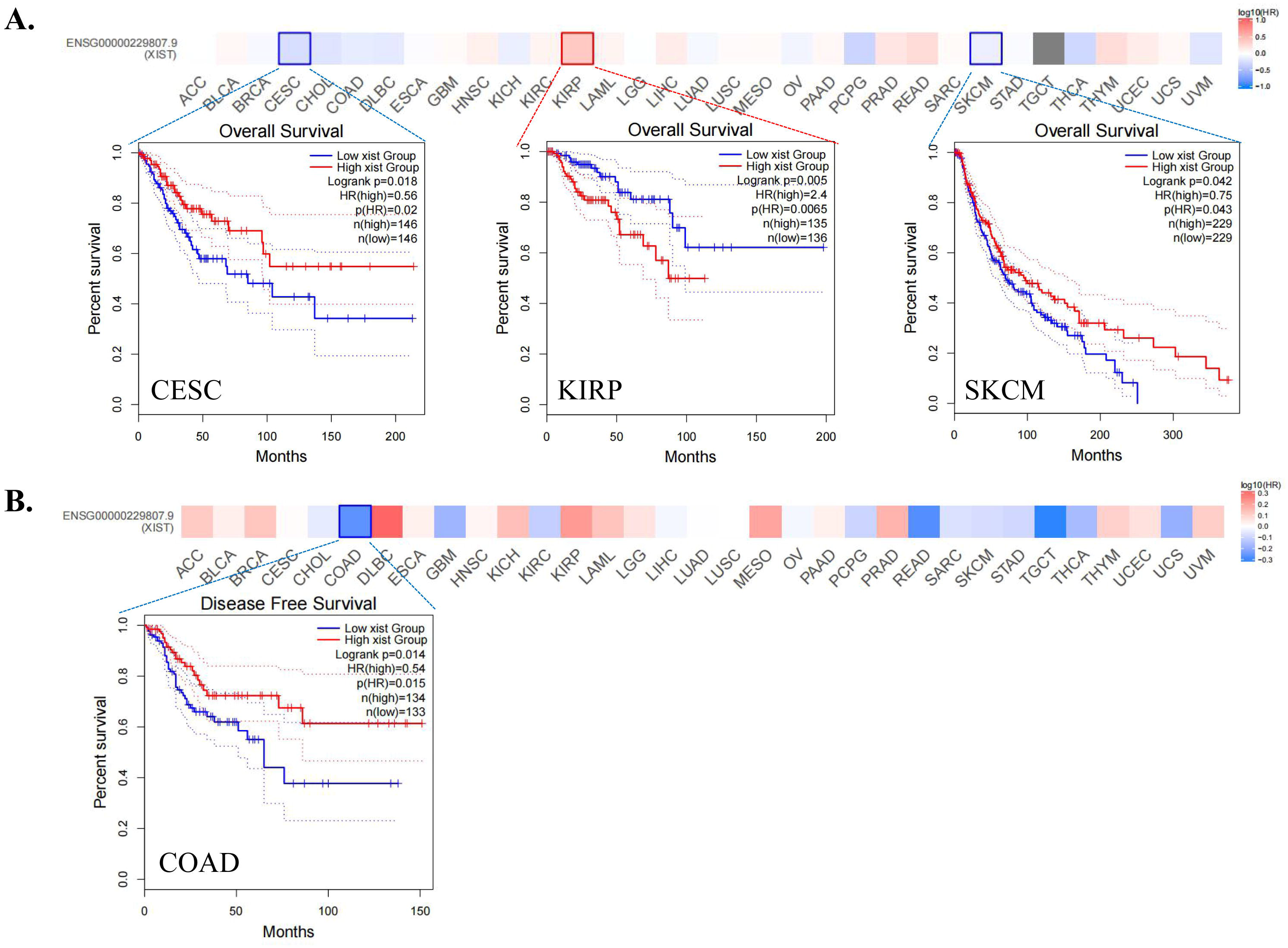
Prognosis analysis of XIST across cancers in GEPIA. **(A)** OS; **(B)** DFS. Red represented high risk and blue represented low risk.

**Figure 3.**
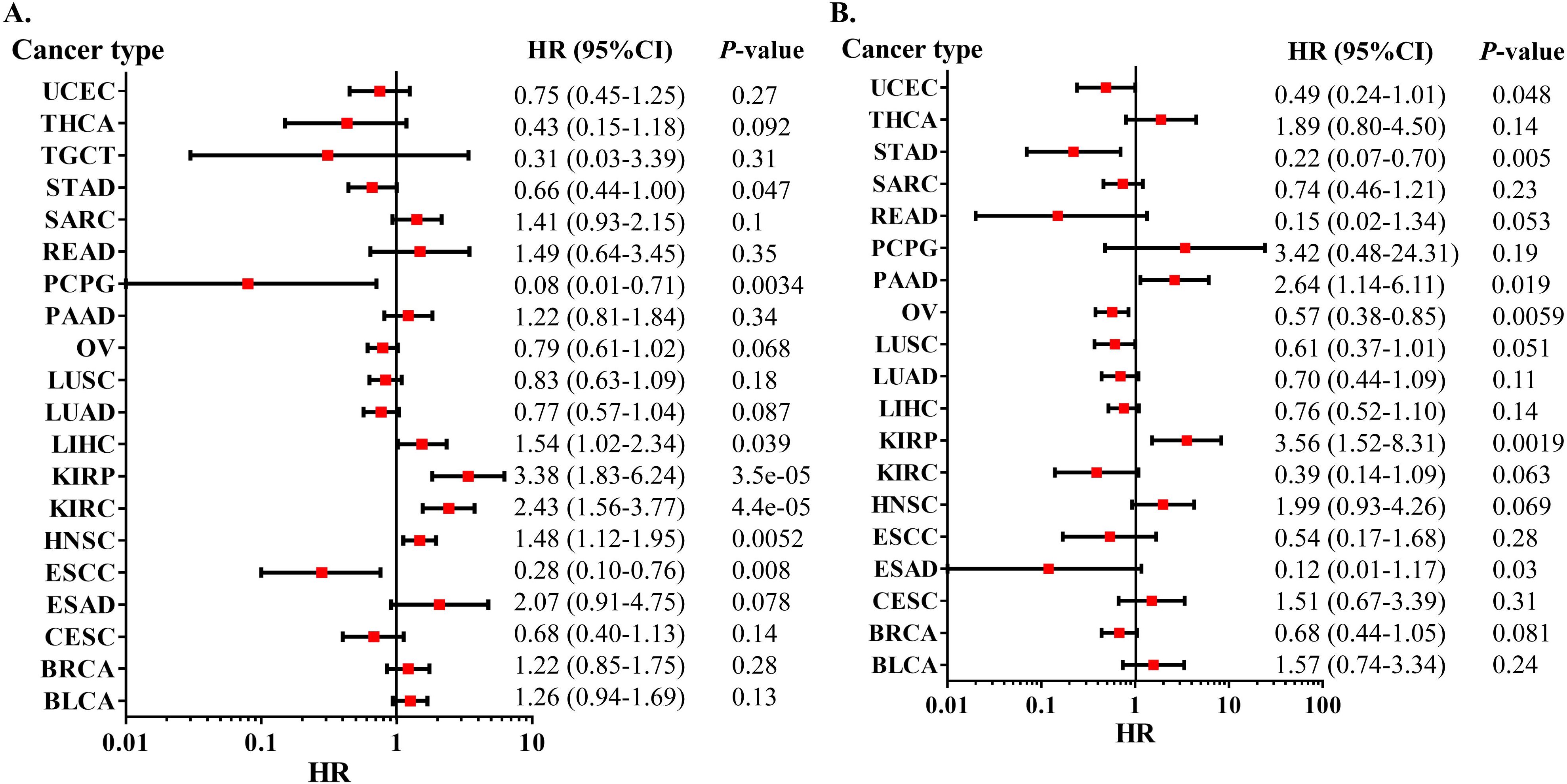
Prognosis analysis of XIST across cancers in Kaplan-Meier Plotter. **(A)** OS; **(B)** RFS.

**Figure 4.**
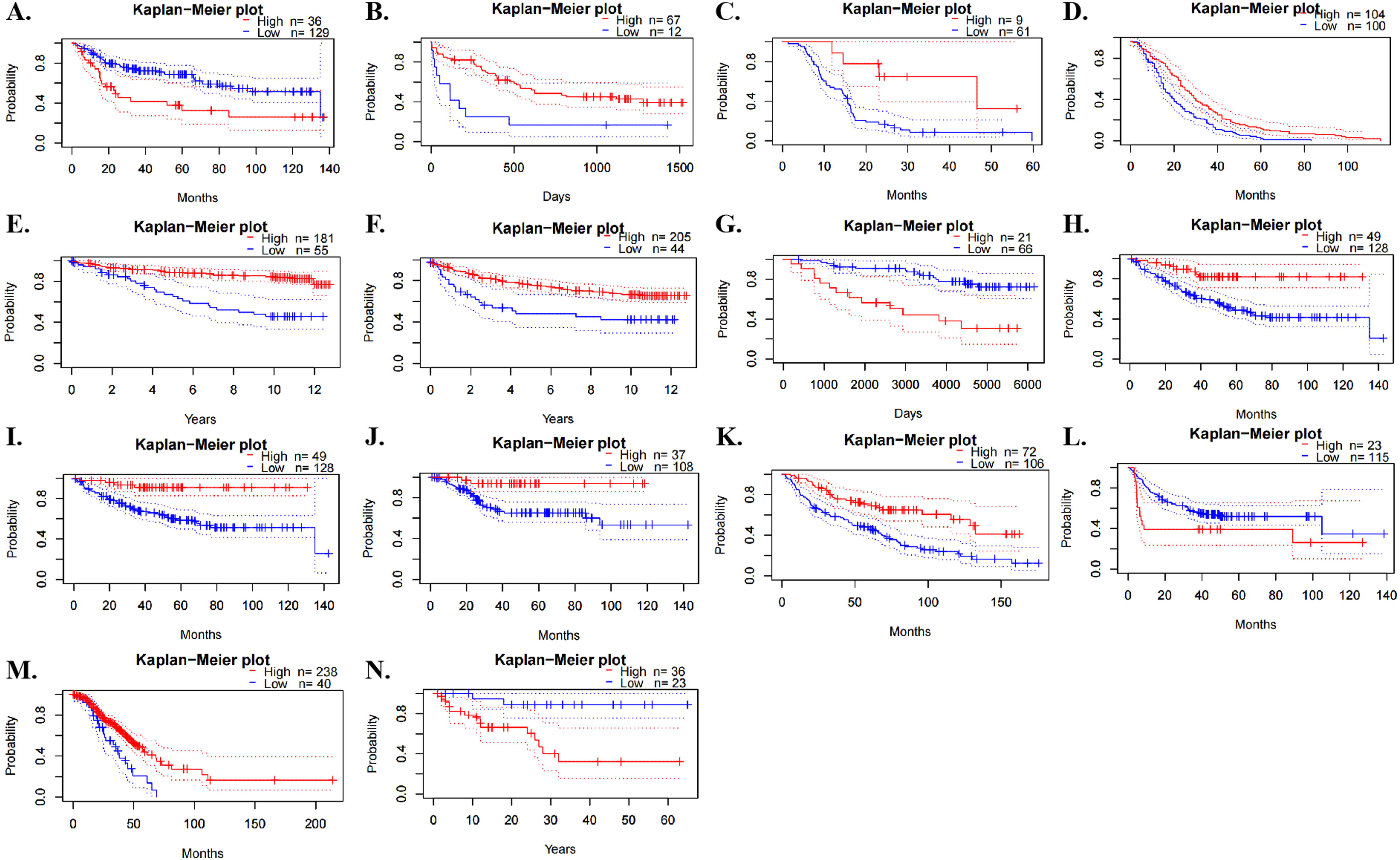
Prognosis analysis of XIST across cancers in PrognoScan. **(A)** OS of BLCA; **(B)** OS of AML; **(C)** OS of GBM; **(D)** RFS of BRCA; **(E)** DSS of BRCA; **(F)** DFS of BRCA; **(G)** DMFS of BRCA; **(H)** OS of COAD; **(I)** DSS of COAD; **(J)** DFS of COAD; **(K)** OS of LUAD; **(L)** RFS of NSCLC; **(M)** OS of OV; **(N)** OS of Renal cell carcinoma.

### Interactions of XIST with proteins and miRNAs across cancers

StarBase discovered that XIST could potentially interact with 29 proteins (Figure 5A) and 191 miRNAs (Figure 5B) across cancers. Over 8 proteins were significantly related to XIST in 4 types of cancers, including BRCA, KIRC, OV and UCEC (Figure 5A). Among them, TNRC6, DGCR8, C17ORF85, ZC3H7B, SFRS1 and TIA1, were obviously positively associated with XIST in ≥ 3 cancers; while eIF4AIII, FXR1, FXR2 and C22ORF28 were significantly negatively correlated with XIST in ≥ 3 cancers (Figure 5A). Although these miRNAs might be sponged by XIST according to the prediction of LncRNA - miRNA interactions from starBase, their expression levels were different among tumors (Figure 5B). Over 10 miRNAs were negatively related to XIST in KIRC, LAML, COAD and READ, while over 10 miRNAs were positively associated with XIST in KICH, LUAD and OV (Figure 5B). In addition, there were more than a dozen miRNAs both negatively and positively correlated with XIST (≥ 10, respectively) in BLCA, BRCA, SKCM and UCEC (Figure 5B). Then, we selected out 4 miRNAs, including miR-103a-3p, miR-107, miR-130b-3p and miR-96-5p, which were all negatively related to XIST expression in more than 3 types of cancers. Furthermore, 128 ceRNAs were found to compete with XIST for miRNAs binding and were positively related to XIST across cancers, especially BRCA and UCEC (Figure 6A). Particularly, expressions of ZFX and TXLNG were both positively correlated with XIST in over 10 cancers. According to TargetScan, miRBase and starBase, we scheduled a network of LncRNA - miRNAs - mRNAs (Figure 6B).

**Figure 5.**
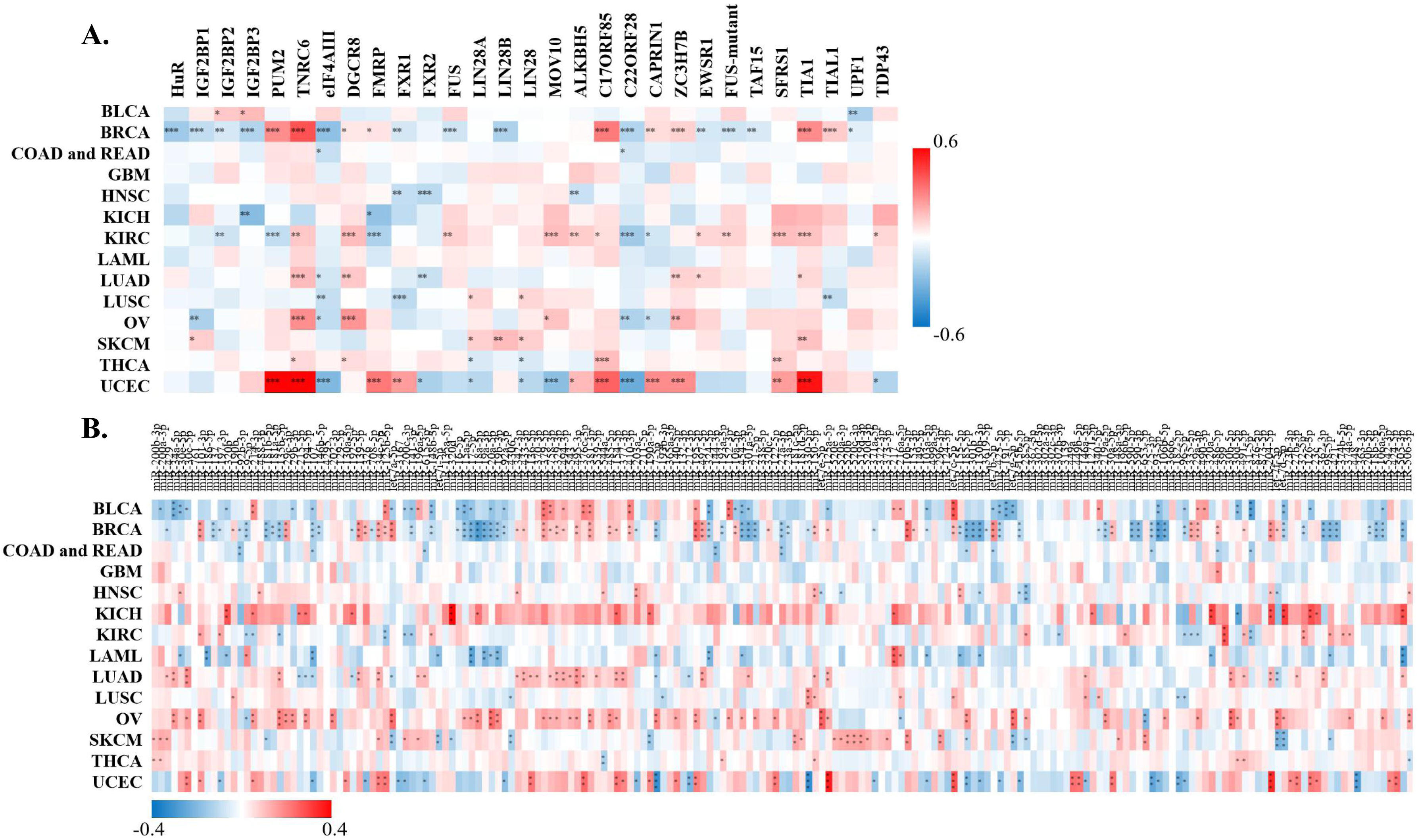
Correlation of XIST with predicted XIST-interacting proteins and miRNAs in starBase. **(A)** Predicted XIST-interacting proteins; **(B)** Predicted XIST-interacting miRNAs. Red represented positive correlation and blue represented negative correlation. * *P* < 0.05, ** *P* < 0.01, *** *P* < 0.001.

**Figure 6.**
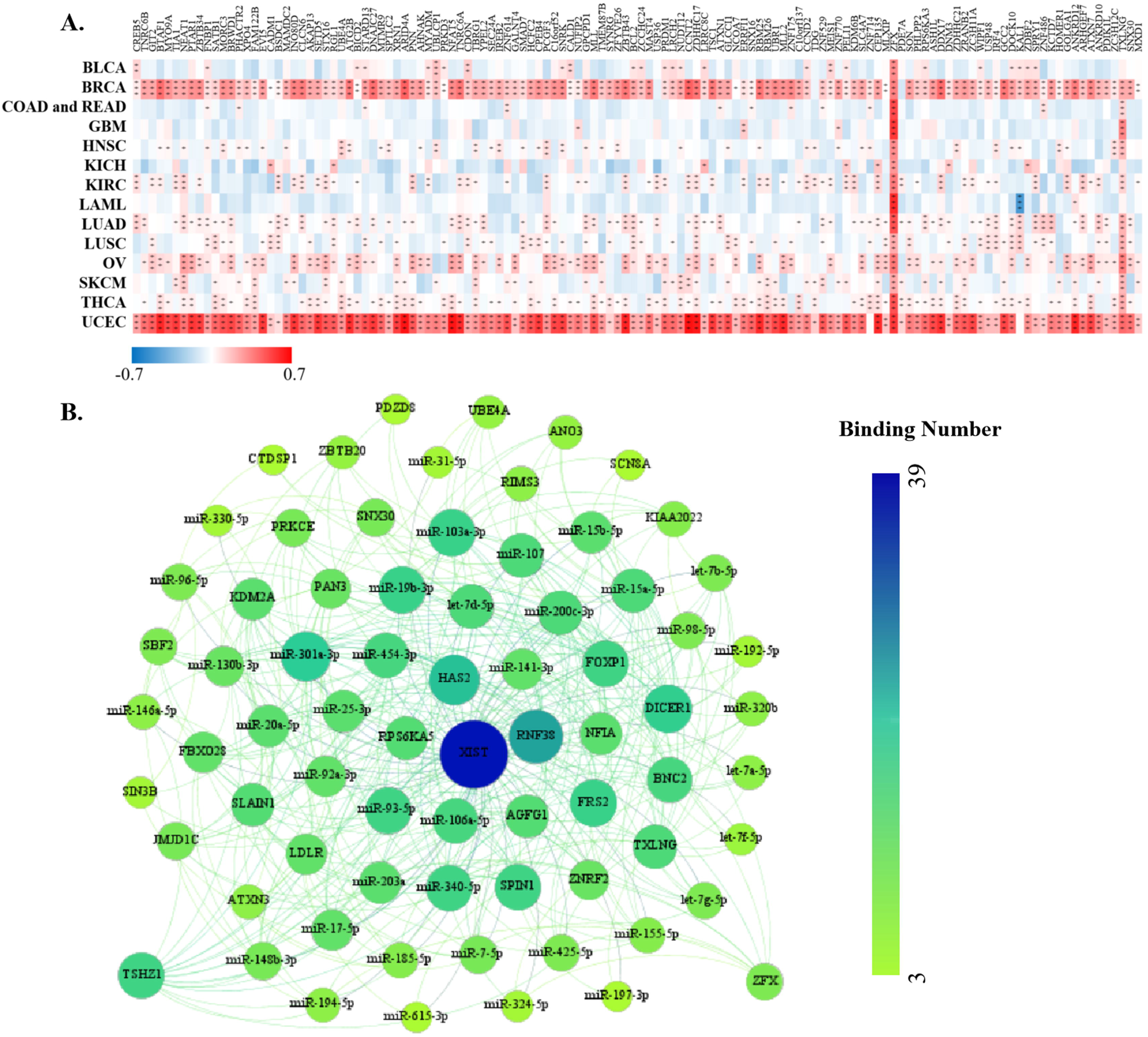
Correlation of XIST with ceRNAs in starBase. **(A)** ceRNAs. Red represented positive correlation and blue represented negative correlation. * *P* < 0.05, ** *P* < 0.01, *** *P* < 0.001. **(B)** A network of XIST/ceRNAs - miRNAs - mRNAs.

### Correlation between XIST and immune infiltration levels in multiple cancers

To determine the role of XIST in TIME, we analyzed the correlation of XIST expression with immune infiltration through TIMER 2.0 database. Its expression was positively related to infiltration levels of T cell CD4+ memory resting, T cell CD4+ memory, T cell CD4+, Tregs and mast cell in over 8 types of cancers, while it was negatively correlated with T cell CD4+ Th1, Macrophage and T cell NK in more than 8 types (Figure 7A). Hence, XIST might influence immune infiltration in the TME.

**Figure 7.**
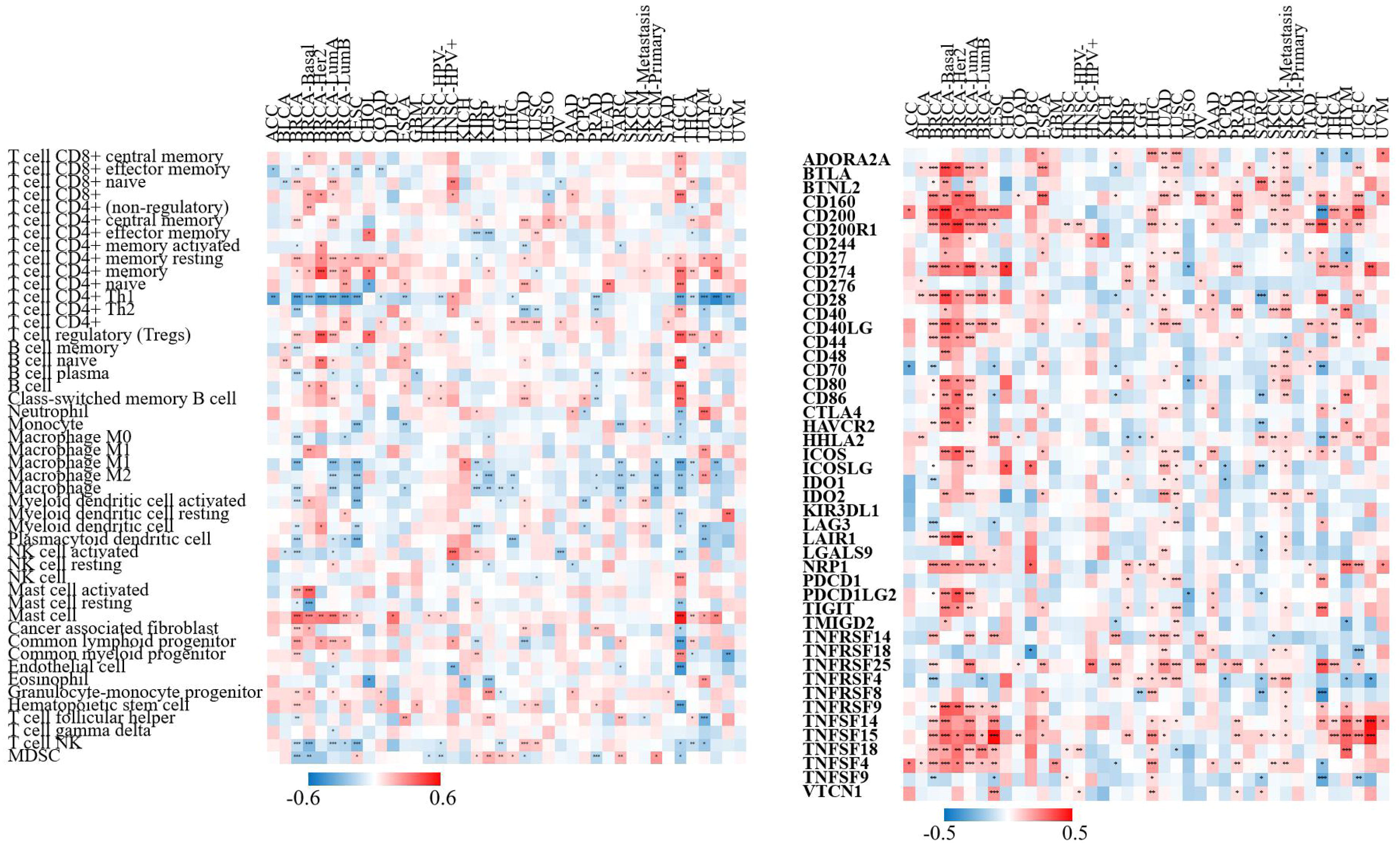
The correlation of XIST with immune infiltration levels and checkpoint markers across cancers in TIMER2. **(A)** immune infiltration levels; **(B)** immune checkpoint markers. Red represented positive correlation and blue represented negative correlation. * *P* < 0.05, ** *P* < 0.01, *** *P* < 0.001.

### Correlation between XIST and immune checkpoint markers in multiple cancers

To investigate the immunoregulatory mechanism of XIST, we investigated more than 40 common immune checkpoint markers in TIMER 2.0 across cancers. Generally, XIST expression was positively related to about half of these markers in various cancers, such as CD2000, CD200R1, CD274, members of the tumor necrosis factor (TNF) ligand family, and so on (Figure 7B). Figure 8 showed the correlation of XIST expression with 35 immune cell types. Notably, its expression was positively related to most of these cell types (> 20) in BRCA. In addition, XIST expression was also positively associated with over 10 cell types in CESC, ESCA, LIHC, LUAD, LUSC, PAAD, PRAD, SKCM-Metastasis, STAD, TGCT and THYM. However, XIST expression was negatively linked with some immune cell types in several cancers, including KIRC (7 cell types), SARC (5 cell types) and TGCT (5 cell types). These results implied that abnormal expression of XIST might become a vital factor for survival of cancer patients through its regulation on the TIME.

**Figure 8.**
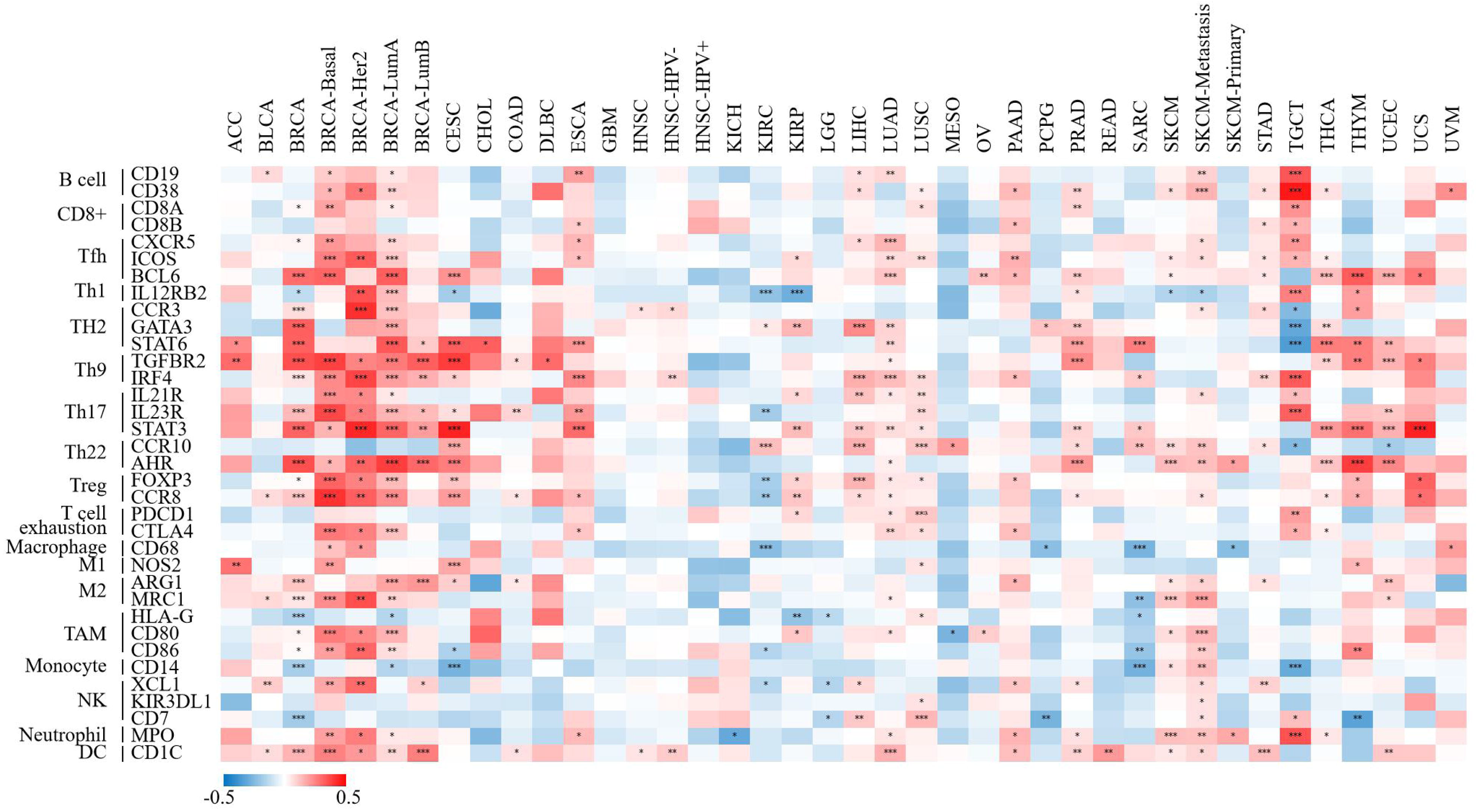
The correlation of XIST with immune cell types in multiple cancers in TIMER2. Red represented positive correlation and blue represented negative correlation. * *P* < 0.05, ** *P* < 0.01, *** *P* < 0.001.

### Correlation between XIST and representative gene mutation across cancers

Next, we analyzed the relationship of XIST expression with 47 types of tumor-associated gene mutation to further explore the mechanism of carcinogenesis of dysregulation of XIST. Generally, theses mutations were weakly related to only several types of cancers (< 5 cancer types for each gene mutation, Figure 9). However, in PRAD, XIST expression was significantly negatively associated with mutations of FAT1 and RB1. In READ, its expression was negatively related to 6 gene mutations, including BRCA1, BRIP1, FANCA, NTRK3, PALB2 and TSC1. In addition, XIST was slightly negatively associated with 5 gene mutations in BRCA (APC, ASXL1, BRCA2, ERBB3 and TP53), but positively related to mutation of PIK3CA. These findings reflected that XIST expression might be affected by tumor-associated gene mutation in several cancers, especially BRCA and READ.

**Figure 9.**
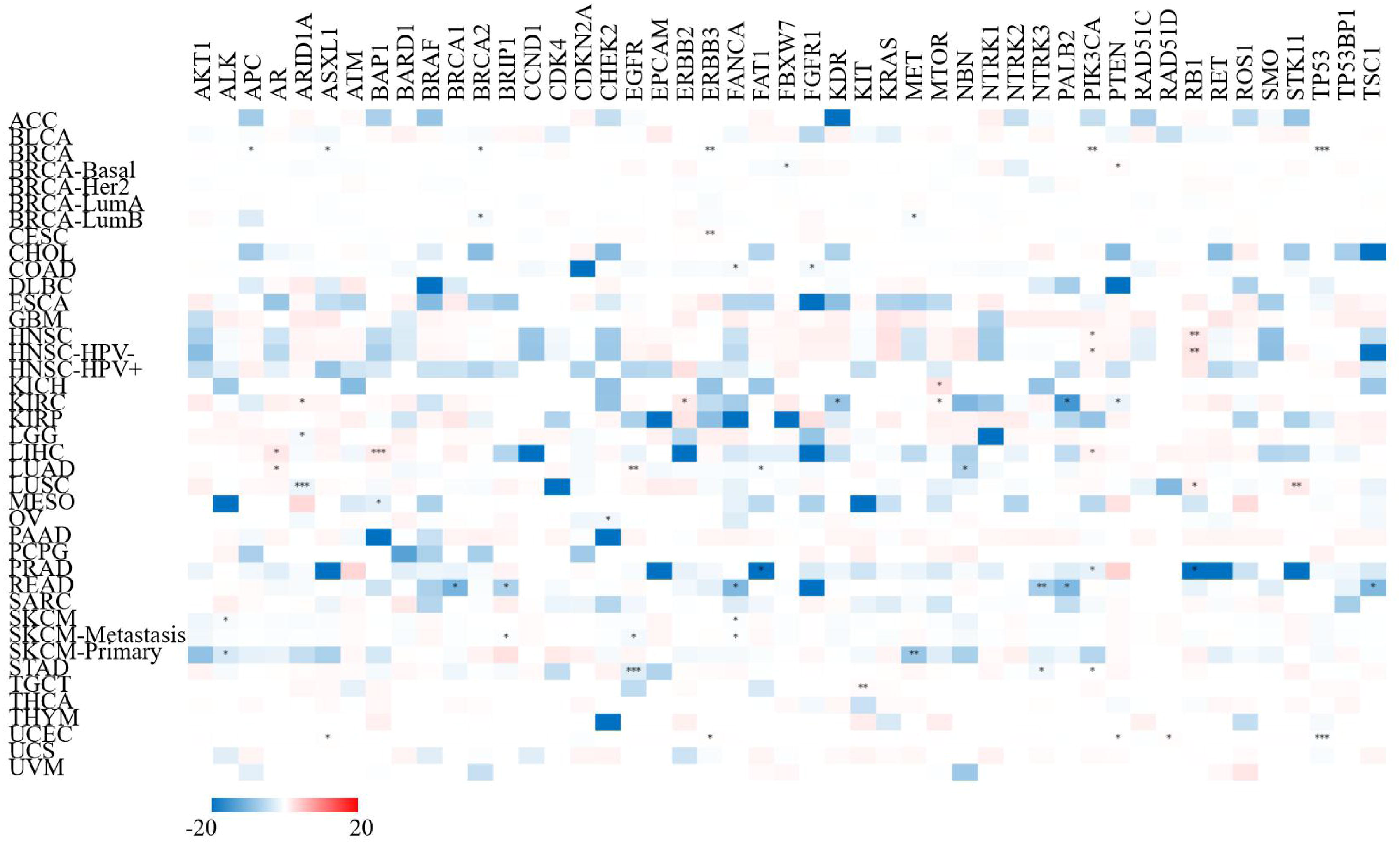
The correlation of XIST with mutations of representative tumor-associated genes across cancers in TIMER2. Red represented positive correlation and blue represented negative correlation. * *P* < 0.05, ** *P* < 0.01, *** *P* < 0.001.

## Discussion

LncRNAs were initially described as mere “transcriptional noise”, but now increasing studies have demonstrated that LncRNAs act as critical modulators in a variety of physiological activities[29]. However, only a relatively limited number of LncRNAs have been confirmed to have critical biological functions, and molecular mechanisms of most LncRNAs have not been illustrated[30]. XIST is a pivotal initiator of imprinted and random X-chromosome inactivation in mammals, and it silences one X chromosome in order to avoid the excessive activation of genes[31]. When dysregulation of XIST occurs, varieties of diseases will emerge due to the escape from X chromosome inactivation[31]. But its mechanisms for pathogenesis are more than that. Recent studies have demonstrated that loss of XIST promotes tumor growth and invasion of BRCA due to the reduction of endogenous competition against onco-miRNAs[16,32]. However, its functions on tumors were not consistent in previous studies. Several researches showed that growth and metastasis of cancer cells were facilitated due to the abnormal upregulation of XIST[17,18,33]. This inconformity resulted in failure to determine whether XIST could become a strong prognostic indicator of cancer patients.

From three databases, Oncomine, TIMER and GEPIA, we found relative expression levels of XIST were different among multiple cancers. Particularly, XIST was down-regulated and predicted a good prognosis in BRCA, COAD and OV, but its expression was inconsistent with outcomes in other cancers. Therefore, we demonstrated that XIST might become a good molecular biological indicator for prognosis of patients with BRCA, COAD and OV. However, several studies showed that XIST was up-regulated in these tumors and was negatively linked with survival rates of patients[34–36]. Hence, further researches need to base upon more large survey samples in order to verify the prognostic roles of XIST in cancers.

It is well known that LncRNAs can function as sponges for multiple tumor-associated miRNAs and indirectly regulate expression of targeted mRNAs[13]. In this research, 191 miRNAs were found to be possibly interacted with XIST. However, not all of them were negatively related to XIST expression, which might be affected by other ceRNAs or other factors. Only 4 miRNAs, including miR-103a-3p, miR-107, miR-130b-3p and miR-96-5p, were negatively linked with XIST in more than 3 types of cancers. All of these 4 miRNAs played promoting roles in carcinogenesis by targeting mRNAs[37–40]. Therefore, XIST might function as a tumor suppressor LncRNA by sponging these onco-miRNAs.

In addition, LncRNAs played fundamental roles in diverse biological and pathological processes by interacting with specific proteins[3]. To search the proteins combined with XIST, we performed the starBase database and preliminarily determined 29 candidate proteins. Among them, only 6 ones were co-expressed with XIST, including TNRC6, DGCR8, C17ORF85 (NCBP3), ZC3H7B, SFRS1 and TIA1, all of which were involved in the development and progression of diverse tumors[41–46]. Thus, we identified that these six proteins potentially interacted with XIST. However, further studies need to confirm whether these proteins can cooperate with XIST to affect biological behaviors and epigenetic pathways of cancers.

Currently, XIST is emerging as a critical immune-related LncRNA and silences a subset of X-linked immune genes[47]. The escape of XIST-dependent genes induced the genesis and recurrence of tumors through diverse perplexing mechanisms[47]. This present research found that XIST was related to infiltration levels of T cell CD4+, Tregs, mast cell, Th1, macrophage and T cell NK in more than 8 types of cancers. One previous study of early-stage LUAD showed that XIST was positively linked with B cells, dendritic cells, follicular helper T cells, mast cells, T cell CD4+/CD8+ effector memory and eosinophil, but was negatively correlated with macrophage, Th2[48]. In addition, XIST expression was positively related to more than 20 immune checkpoint markers and over 10 immune cell types across cancers. Therefore, we demonstrated that XIST could become a immune-related LncRNA and played a pivotal role in immune-oncology.

Genetic variation is associated with various disease, especially carcinomas[26]. Mutations of tumor-associated genes appear in various carcinomas, including base substitution and frameshift mutation[26]. Cancer is essentially a genetic disease where cells either die or cancerate, when a certain number of genetic mutations accumulate in the cells. Our research also showed that XIST might be regulated by gene mutation (including APC, BRCA1, BRCA2, TP53 and PIK3CA) in several cancers, especially BRCA, PRAD and READ. Mutations of these genes modulate expressions of diverse downstream genes and have become important therapeutic targets for anticancer drug development[49,50]. For instance, PARP inhibitors have been used for cancer patients with BRCA1/BRCA2 mutation. The XIST RNA domain number in BRCA1 breast tumor was associated with chromosomal genetic abnormalities, and XIST might become a predictive biomarker for prognosis of patients with BRCA1 breast tumor[27,51]. But BRCA1 was dispensable for XIST RNA coating of the X chromosome[51]. These findings provided clues on the association between XIST and mutation of tumor-associated genes. Hence, future researches should be conducted to explore possible mechanisms of XIST in multiple cancers.

Admittedly, this research still had some limitations. First, it was difficult to analyze the prognostic value of XIST in sub-types of cancers through public datasets. Secondly, since this study was based on pan-cancer data in patients, we failed to prove all the ideas at the same time. Thirdly, our results lack external validation in other public databases or in vitro and in vivo researches. This theoretical work remains to be verified.

## Conclusion

In conclusion, dysregulation of LncRNA XIST was significantly associated with prognosis, miRNAs, immune cell infiltration and mutations of tumor-associated genes in multiple cancers, especially BRCA and colorectal cancer. XIST may act as a novel biomarker for survival and immunotherapy across cancers in the immediate future.

## Declarations

### Ethics approval and consent to participate

Not applicable.

### Consent for publication

Not applicable.

### Availability of data and materials

All data generated or analyzed during this study are included in this published article.

### Authors’ contributions

Conceptualization: WH, CTS and HNW. Methodology: WH, XJG and QXS. Investigation: WH, CTS and JM. Visualization: WH, CTS and JM. Manuscript draft: WH, CTS and JM. Manuscript review and editing: WH, CTS, JM, HNW and QXS. Supervision: HNW and QXS. All authors read and approved the final manuscript.

## Acknowledgements

Not applicable.

